# Boost-HiC : Computational enhancement of long-range contacts in chromosomal contact maps

**DOI:** 10.1101/471607

**Authors:** L. Carron, J.B. Morlot, Matthys V., A. Lesne, J. Mozziconacci

## Abstract

Genome-wide chromosomal contact maps are widely used to uncover the 3D organisation of genomes. They rely on the collection of millions of contacting pairs of genomic loci. Contact frequencies at short range are usually well measured in experiments, while there is a lot of missing information about long-range contacts.

We propose to use the sparse information contained in raw contact maps to determine high-confidence contact frequency between all pairs of loci. Our algorithmic procedure, *Boost-HiC*, enables the detection of Hi-C patterns such as chromosomal compartments at a resolution that would be otherwise only attainable by sequencing a hundred times deeper the experimental Hi-C library. Boost-HiC can also be used to compare contact maps at an improved resolution.

Boost-HiC is available at https://github.com/LeopoldC/Boost-HiC

## 1 Introduction

Chromosomal conformation capture (3C) has originally been developed to identify sets of DNA segments in close spatial proximity within a cell nucleus, and thus get an experimental in vivo access to the genome 3D organisation. It relies on the chemical fixation of chromosomal contacts, digestion with a restriction enzyme and a subsequent re-ligation of the cross-linked fragments [3]. Next-generation sequencing techniques brought this protocol to the whole-genome scale (Hi-C) in cell populations [14] and an everyday increasing number of data-sets are now available. These datasets are usually difficult to compare since they can largely vary in terms of quality and sequencing depth. The highest resolution that can be achieved is in theory the size of the restriction fragments but very few datasets reach this resolution in practice. As a stunning example of pushing the experiments to its limits, Rao and colleagues provided the first 1kb-resolution contact map of the human genome, obtained by sequencing 4 billions of read pairs [9]. This dataset as been used in several studies since its release and many other data-sets have been produced in other human cell types and other organisms, but generally at a lower resolution [11]. In mouse, another very high resolution data-set has recently been produced, covering the in vitro differentiation of mouse neurons [1]. These maps are often interpreted by determining the position along genomes of 3D structural features such as Topologically Associated Domains (TADs) boundaries, loops and compartments (e.g. in [9]). Many algorithms have been developed to delineate these features of the 3D genome organisation but their power is always hindered by the limited resolution of the dataset itself. While in principle the resolution can be improved arbitrarily by increasing the sequencing depth, this also dramatically increases the financial cost of the experiment. Improving the resolution by computational methods is therefore a good option and a methods relying on deep neural networks (HiCPlus) has recently been developed [18]. In this method, contact enhancement relies on determining more precisely low contact frequencies from the contact patterns of the genomic neighbours. Adopting a different viewpoint, our guideline is to use instead the path-length on the contact graph [7] as a quantitative index indicating how low-confidence contact frequencies are to be reinforced (Figure 1A). The shorter the path on the contact graph between two genomic sites, the higher should be their contact frequency. In this scheme, low confidence long range contacts are determined as a sum of higher confidence contacts. The numerical implementation of this principle, Boost-HiC, is thus expected to dramatically improve the accuracy, reliability and usability of lower-resolution contact maps.

## 2 Methods

### 2.1 Hi-C sequence alignment

We used previously published dataset [1] from mouse Embryonic Stem (ES) cells and Cortical Neurons (CN) available as GSE96107. Hi-C reads were processed using the mm9 reference genome with HiC-Pro [12]. Bowtie2 was used with default pipeline parameter *–very-sensitive -L 30 –score-min L,-0.6,-0.2 –end-to-end –reorder*. We analysed replicates for each dataset at a resolution of 10kb separately before merging them at the end of the HiC-Pro pipeline. This merged array defines the raw contact map, *M*.

### 2.2 Hi-C Filtering

Hi-C maps were filtered in two steps. First, empty bins are removed. In a second step, the distribution of the sum of contacts for each bin is fitted using a Gaussian Kernel Density function [8] with parameter *kernel=‘gaussian’, bandwidth=2000*. Since Gaussian Kernel Density function returns the logarithm of the distribution, we took the exponential of the output. The resulting distribution is then used to identify bins for which the sum of contacts has a value below 5 % and above 95 % of the mean contact number. These bins are removed from the contact map. Filtered datasets were finally normalised using the Sequential Component Normalization [2] so that the L1-norm of each column/line is equal to 1.

### 2.3 Contact Probability

Contact probability *P*(*s*) as a function of the genomic distance *s* is computed as the mean number of contacts along each secondary diagonal, where s is its distance to the main diagonal. Since there are less contacts between bins which are more distant than between closer ones, contacts are gathered over bins with a size increasing according to a geometric progression (log-binned) of scale factor 1.01. When necessary, we denote *P_C_*(*s*) the contact probability curve associated with the contact map *C*.

### 2.4 Boost-HiC Implementation

Boost-HiC algorithm operates in two steps. The normalised contact map *C* is first transformed element-wise in a distance matrix (*d*°) as described in [6] according to the formula :

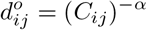

A shortest-path algorithm [4, 15] is then run on *d*° to fill the distance matrix *d*, by defining the distance *d_ij_* between any pair of loci *i* and *j* as the length of the shortest-path connecting them on the weighted contact graph [7]. This complete distance matrix *d* is then turned element-wise into a filled contact map *F*° according to the formula :

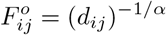

The number of rewired contacts (*nr*) is computed from *C* and *F*° by counting the number of elements (*ij*) taking different values in these two maps. We started with a small value (e.g. 0.05) of the exponent α and then increased this value by step of 0.01 and computed *nr*. When *nr* started to increase, we stopped the procedure and kept this value for the exponent *α*.

We adjusted the contact probability curve associated with *F*° so that it matches exactly the one computed for *C*. Each contact 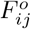 for sites *i* and *j* separated by a genomic distance *s* = |*i* -*j* | is adjusted by the ratio between the raw probability against the improved one :

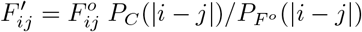

The code recapitulating Boost-HiC steps for a given value of alpha is available at https://github.com/LeopoldC/Boost-HiC.

### 2.5 HiCPlus Implementation

HiCplus as been used as described in [18]. We modified the original code in order to apply the algorithm of the whole contact map whereas it was originally developped to enhance short range contacts only. The version of the code we used is available at https://github.com/jbmorlot/HiCPlus/.

### 2.6 Downsampling

In order to quantify the efficiency of our algorithm, we wanted to see if it is capable of inferring the same information as a deeply sequenced map when starting from a map with less contacts. To implement this test, we sampled the raw map with a binomial probability. For every couple of bins *i* and *j* in the raw map, the number of contacts *M_ij_* between these two genomic regions is replaced with a random number generated from a binomial distribution of parameters *M_ij_* and *k*. We downsampled the matrix with 3 different values of *k* : 0.1, 0.01 and 0.001 corresponding respectively to the downsampling of the whole map at 10 %, 1 % and 0.1 %.

### 2.7 Compartments and TADs

Compartments were obtained as in [14]. First, normalised contact maps were transformed to their observed/expected ratio. Correlation matrices of these ratios, also termed correlation maps, were then computed. The sign of the first eigenvector of these correlation maps was used to infer two compartments. A (active) and B (inactive) compartments were then identified by computing the gene density in each compartment and assigning the label A to the highest-density compartment.

**Figure 1:**
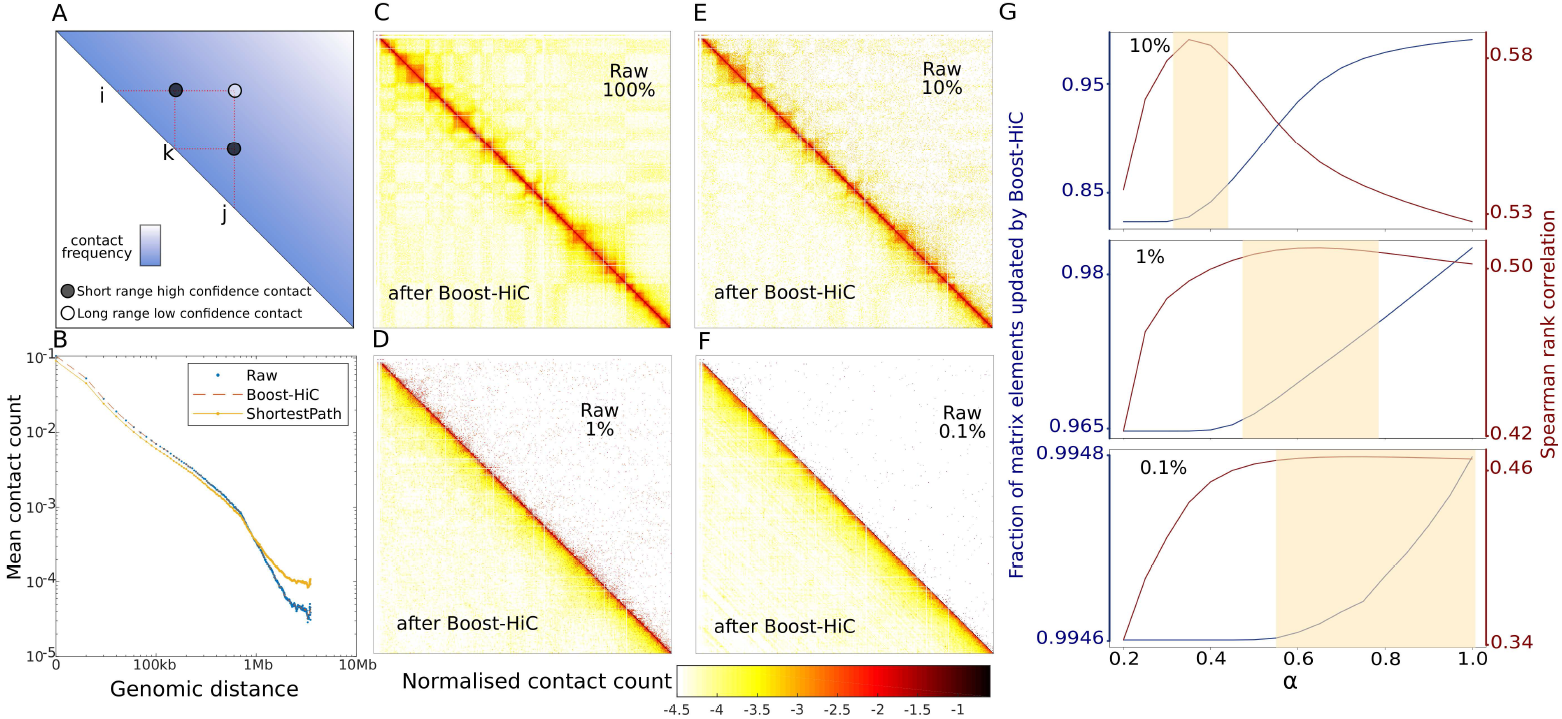
**A)** Schematic explanation of our shortest-path enhancement algorithm on Hi-C map : the low-confidence long-range contact between sites a and b (white dot) is enhanced using the information on shorter-range higher-confidence contacts (black dots) with intermediary nodes (here c). On the contact graph, the shortest-path between a and b goes through *C* and the enhanced contact value is related to its length. **B)** Contact probability curve *P*(*s*) of ES-cells from their Hi-C map downsampled at 10 % of contacts. The blue points display the original probability of the downsampled map. Yellow line displays the probability after information enhancement by means of our shortest-path method. The red line displays the improved contact probability at the end of Boost-HiC algorithm. **C)** to **F)** panels display the mouse ES-cell contact map (for a region on chromosome 16 located between from 100kb and 29,8Mb) at different downsampling rates (in order) : 100 %, 10 %, 1 % and 0.1 %. Upper parts displays the raw contact map at fixed downsampling. Lower parts shows the map improvement after implementation of Boost-HiC. Colormap in log10 scale [2].**G)** Pearson correlation (in red) between full raw maps and boosted maps for different downsampling rates and value of the parameter *α* used in the Boost-HiC algorithm. The blue line shows the fraction of rewired contacts between the downsampled map and the boosted map. From top to bottom, we compared the 100 % map with respectively the 10 %, 1 % and 0.1 % map enhanced with Boost-Hi-C.

To infer the position of Topologically Associating Domains (TADs), we computed a “TAD border score” for each 10 kb position *p* along the genome. This score obtained by summing the contact map elements over a square region of size *h* with its corner located on the main diagonal, at the poistion *p*. We next plotted the score along the genome by sliding the square along the diagonal of the contact map [5]. We chose here a size *h* of 300kb. Local minima of this score determined the TAD borders [13].

## 3 Results and Discussion

### 3.1 Boost-HiC partially restores the resolution of downsampled contact maps

Hi-C datasets are generally processed in the form of contact maps, in which the contact frequency between two loci is determined by normalising the number of times they have been found together in the Hi-C library. At fine resolution (i.e. when contacts are aggregated over small genomic bins), this number is frequently equal to zero due to the finite sequencing depth. The issue is currently circumvented by working at a lower resolution, by binning the restriction fragments into larger and less numerous regions. We propose a computational alternative, Boost-HiC, to infer the missing fine-resolution contact frequencies from the knowledge of the measured contacts, and get a complete fine-resolution contact map. The procedure is based on the computation of shortest-paths between any pair of genomic loci on the contact network, a method that we introduced previously for 3D reconstruction [6, 7]. More precisely, we first compute the dependency of the contact probability on a normalized contact map [2] *C* with respect to the genomic distance *s* between two loci, *P*(*s*). We also transform the contact map, with elements *C_ij_*, in a distance map with elements

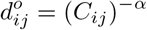

where the exponent *α* is a parameter that will be later optimised. This initial distance map *d*° defines a weighted contact network, where the link between the loci *i* and *j* is given a length 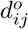. We then compute an updated distance map using a shortest-path algorithm : the distance *d*1_ij_ between any two loci *i* and *j* is set to the minimal distance on the graph between nodes *i* and *j*, that is, the length of the shortest path relating*i*and *j*. In this way we get a full distance map *d*1 with finite elements *d_ij_*, even when the measured contact frequency between the loci *i*and *j* vanishes. This distance map is then converted into a contact frequency map *F* ° by inverting the above relation element-wise, namely 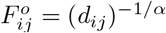. Since short-range contact values were often used to infer longer-range contact values, the *P*(*s*) curve may change after the procedure. A proper element-wise rescaling of *F*° restores the original contact probability curve *P*(*s*) at this step (Figure 1B). The resulting map *F’* is finally normalised using the Sequential Component Normalization procedure [2] to give the final ‘boosted’ contact map *F*. This map *F* lies at the same fine resolution than the original one, but vanishing elements are replaced with their inferred values.

To assess the capabilities of our algorithm, we generated 3 downsampled contact maps sampling 10 %, 1 % and 0.1 % of the initial contacts from a high-resolution map of region of chromosome 16 in mouse at 10kb resolution (see Methods). We then constructed the corresponding boosted maps (Figure 1 C-F). In order to quantitatively compare the boosted map and our objective (i.e. the original high resolution version), we computed the Spearman correlation between the boosted maps and the high-resolution map for different values of our free parameter *α*. While some other methods have been developed to compare contact maps from different origin [10,16,17], this straightforward measure works well when comparing downsampled maps with the original one. We found that the downsampled maps displayed a reduced correlation of 0.51, 0.26 and 0.11 for 10 %, 1 % and 0.1 % downsampling respectively. When *α* is optimally chosen, this correlation increases to 0.59, 0.51, 0.47 for the boosted maps obtained for the corresponding downsampled maps. In comparison, HiCPlus [18], which uses a deep-learning approach where contacts are enhanced using the information contained in the contacts established between the adjacent sites along the genome, achieves correlation values of 0.59, 0.44 and 0.34. This latter result could be explained by the fact that the enhancing procedure HiCPlus was specifically designed for short-range contacts whereas Boost-HiC mostly improves low, long-range contact frequencies.

### 3.2 Optimising the parameter *α*

The different values of the Spearman correlation obtained for increasing values of our free parameter *α* showed that unless *α* is chosen below 0.1 or above 1, the choice of its value has a mild effect on the reconstruction efficiency (e.g. it changes from 0.53 to 0.58 for a down-sampling of 10 %, Figure 1 E)). The best *α* (i.e. yielding to the most accurate reconstruction of the full initial map) is equal to 0.25, 0.5 and 0.6 for different downsampling rates and it is a priori difficult to know which value to use in a practical case. For low *α* values, only vanishing elements of the sparse original matrix are changed into non-zero values. When *α* increases, the number of elements of the matrix that are updated in the Boost-HiC procedure increases and non-zero elements are also modified. In order to see whether the optimal value for *α* depends on the number of modified values, we plotted the number of non-zeros values that are reassigned as a function of the downsampling rate and *α* value. As expected, as the downsampling rate increases, more and more elements become equal to zero in the downsampled matrix. Interestingly, we showed that for all three downsampling rates, the optimal value of *α* corresponds to the updating of vanishing (i.e. all the zeros and few non-zeros) elements only (Figure 1 F). Importantly, this can give us a way to choose an optimal *α* value in real cases, where we have only access to sparse data. We therefore implemented a search of the optimal *α*, which corresponds to a re-assignment of all the zeros and almost 10 % of the non-zeros elements only.

Since the alpha optimisation step can be long for large matrices, we propose in this case to estimate the optimal alpha only based on the sparsity of the matrix. For low sparsity data (i.e. lower than 0.9) an alpha value of 0.25 works well whereas for high sparsity data (i.e. higher that 0.99) alpha should be chosen close to 0.6. For intermediate sparsity, alpha should ideally be taken between those two bounds.

### 3.3 Boost-HiC enables the precise determination of compartments from low resolution maps

As the contact patterns vanish in down-sampled maps (Figure 1 C-F), the detection of A/B compartments using the state-of-the-art procedure is gradually impaired. At 0.1 % down sampling compartments cannot be identified anymore (Figure 2 A). When the Boost-HiC procedure is applied to the down-sampled maps, the detection of these structural features is restored, showing the ability of our computational strategy to reliably detect 3D features from sparse data (Figure 2 B). Using HiCPlus to enhance the down-sampled maps also resulted in a partial recovery of the compartments (Figure 2 C). All these results are summarised in Figure 2 D), which shows the bins that are attributed to the good compartment, as determined from the high resolution map, for different down-sampling values and enhancement methods. Boost Hi-C robustly allows a better prediction of compartments from low resolution maps.

On the other hand, since TADs are determined from short range contacts, their detection in not changed by the Boost-HiC procedure (Figure 2 E). Indeed the TAD border signal is not (or very mildly) modified by the Boost Hi-C procedure. On the other hand, we noted that HiCPlus does change the signal around the diagonal, although the position of TADs borders as minima of the signal are not modified.

### 3.4 Boost Hi-C enables high-resolution comparison between contact maps

We finally assessed the performance of the Boost-HiC procedure for comparing contact maps obtained in different biological conditions. The comparison is usually done by computing the element-wise log-ratio of two such maps. In order to be informative, this comparison can only be done at a resolution for which the signal-to-noise ratio is high. On Figure 3, we compare two contacts maps from two different cell types at different sequencing depths, with and without using Boost-HiC. We compared here mouse Embryonic Stem cells (ESC) with Cortical Neurons (CN). Even if these maps only represent a sub-region of mouse chromosome 16, many pairwise contacts between 10kb bins are equal to zero in a typical Hi-C experiment, even at the highest coverage (Figure 3 A). When computing the log ratio, these elements of the ratio map will therefore be undefined, as they will be equal to +/− infinity (in green and yellow on Figure 3, right panels). In contrast, when the maps have been complemented with Boost-HiC prior to this differential analysis, these elements will always be filled with a finite value (in red and blue on Figure 3, right panels). As can be expected, this effect is even more pronounced at low resolution (e.g. at 10 % downsampling in Figure 3B)

**Figure 2:**
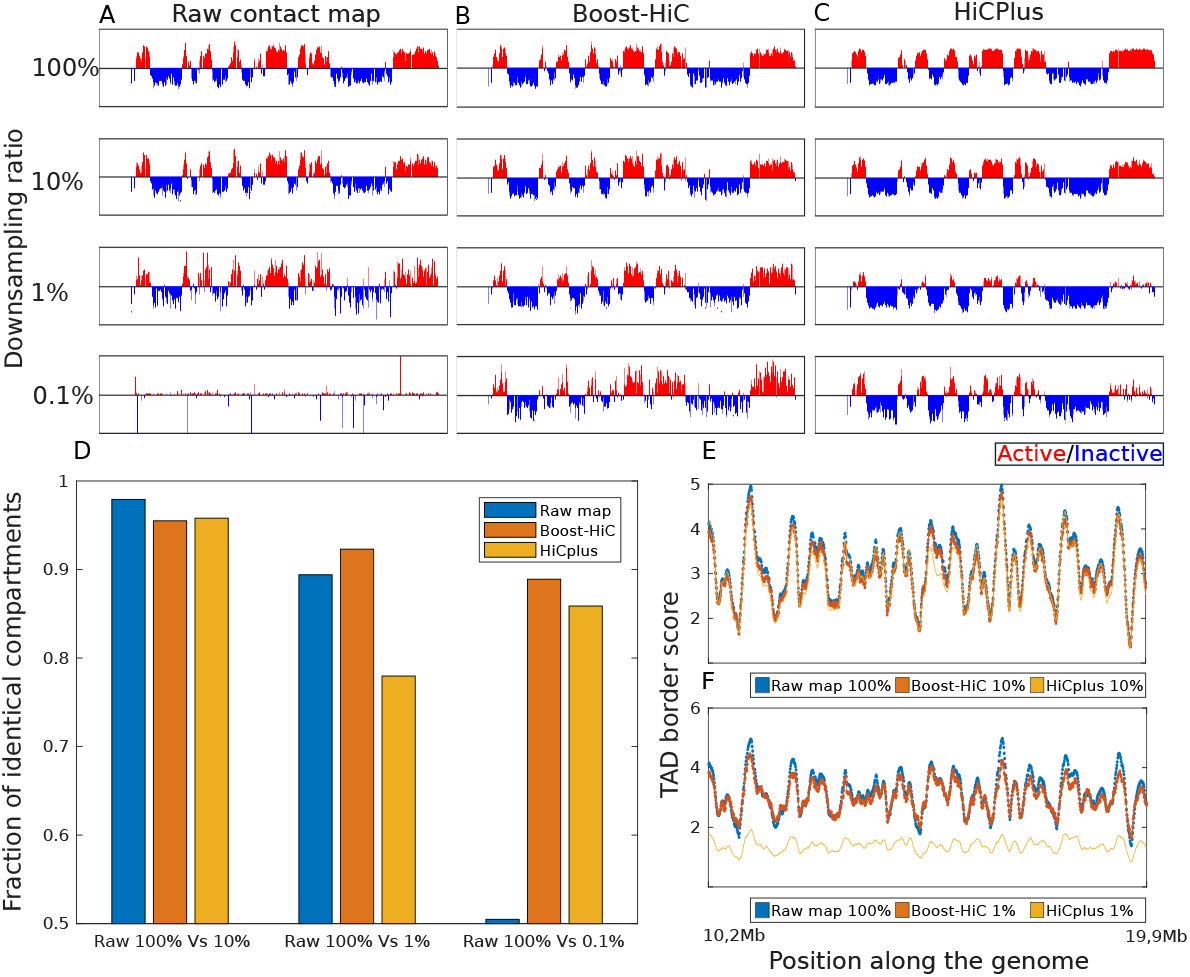
**A)** Barplot of the first eigenvector of mouse ES-cell correlation map (for a region on chromosome 16 located between 100kb and 29,8Mb) at different downsampling rates (in order from top to bottom) : 100 %, 10 %, 1 %, 0.1 %. B) Similar barplot computed after application of the Boost-HiC algorithm. **C)** Similar barplot computed after application of HiCPlus algorithm [18] **D)** Fraction of regions attributed to the correct compartment from downsampled maps using different enrichment methods for 10 %, 1 % and 0.1 % downsampling. **E)** TAD border signal along the genome coordinate (from 100kb to 29,8Mb of chr16). In all plots the blue line shows the score for the raw normalised dataset at 100%. In the top plot red and yellow line correspond to the dataset downsampled at 10% enhanced respectively with Boost-HiC and HiCPlus methods. **F)** Same plot as **E)** with Boost-HiC and HiCPlus methods used on the dataset downsampled at 1%.

**Figure 3:**
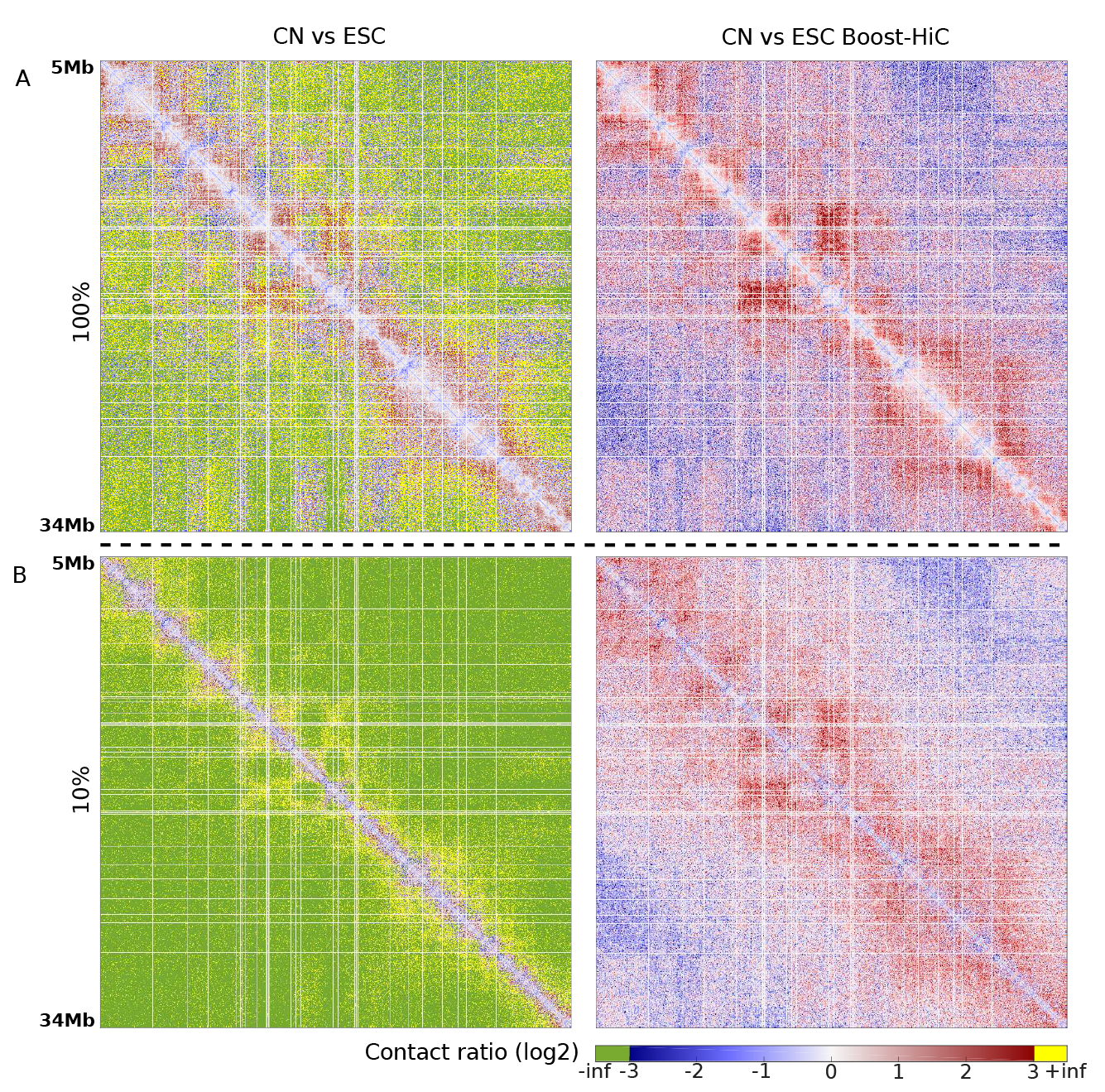
Log2-ratio between CN and ES-cell contact maps, before (left panels) and after (right panels) application of the Boost-HiC method. An null value in one dataset will give an infinite value in the log_2_-ratio. **A)** Log_2_-ratio for maps without downsampling. **B)** Log_2_-ratio for maps downsampled at 10 %.

## 4 Conclusion

We present here Boost-HiC, a computational method aiming at enhancing low resolution chromosomal contact maps. We showed that the resulting maps can be efficiently used in order to recover chromosomal compartments at high resolution. We also show that the procedure can also be efficiently used when comparing Hi-C contact maps when looking for differential contacts between two conditions.

